# Pulmonary Matrix Derived Hydrogels from Patients with Idiopathic Pulmonary Fibrosis Induce a Proinflammatory State in Lung Fibroblasts *In Vitro*

**DOI:** 10.1101/2023.05.03.539323

**Authors:** JG Fernandez Davila, DW Moore, J Kim, JA Khan, AK Singh, M Lemma, CS King, SD Nathan, LR Rodriguez, GM Grant, JL Moran

**Affiliations:** Department of Biology, George Mason University, 10900 University Blvd., Manassas, VA 20110, USA; Department of Mechanical Engineering, George Mason University, 10920 George Mason Circle, Manassas, VA 20110, USA; Department of Bioengineering, George Mason University, 10920 George Mason Circle, Manassas, VA 20110, USA; Inova Advanced Lung Disease and Transplant Program, Inova Fairfax Hospital, 3300 Gallows Road, Falls Church, VA 22042, USA

**Keywords:** Pulmonary Fibrosis, Decellularized Matrix, Mechanobiology, Three-dimensional, Culture, Fibroblast-Immune Crosstalk

## Abstract

Idiopathic pulmonary fibrosis (IPF), one of the most common forms of interstitial lung disease, is a poorly understood, chronic, and often fatal fibroproliferative condition with only two FDA-approved medications. Understanding the pathobiology of the fibroblast in IPF is critical to evaluating and discovering novel therapeutics. Unfortunately, our ability to interrogate this biology *in vitro* is greatly limited by the well-documented effects of tissue culture plastic on the fibroblast phenotype. Using a decellularized lung matrix derived from IPF patients, we generate three-dimensional (3D) hydrogels as *in vitro* models of lung physiology and characterize the phenotype of fibroblasts seeded into the hydrogels. When cultured in our hydrogels, IPF fibroblasts display differential contractility compared to their normal counterparts, lose the classical myofibroblast marker α-smooth muscle actin, and increase expression of proinflammatory cytokines compared to fibroblasts seeded two-dimensionally (2D) on tissue culture dishes. We validate this proinflammatory state in fibroblast conditioned media studies with monocytes and monocyte-derived macrophages. These findings add to a growing understanding of the lung microenvironment effect on fibroblast phenotypes, shed light on the potential role of fibroblasts as immune signaling hubs during lung fibrosis, and suggest intervention in fibroblast-immune cell crosstalk as a possible novel therapeutic avenue.

## 1. INTRODUCTION

Idiopathic pulmonary fibrosis (IPF) is a chronic fibroproliferative disease characterized by dysregulated wound healing and severe scarring in the lung(Betensley *et al*., 2016; Sgalla *et al*., 2018). Under normal conditions, wound healing proceeds at the molecular level via a tightly regulated sequential cycle of inflammation, proliferation, and remodeling. When this process is dysregulated, wound healing proceeds indefinitely, resulting in progressive scarring, tissue stiffening, and physiological restriction(Smithmyer *et al*., 2014; Moretti *et al*., 2022). The remodeling of the pulmonary extracellular matrix (ECM), orchestrated by lung resident fibroblasts, is critical to the progression of fibrotic diseases such as IPF. Fibroblasts are the primary contributors to the extensive deposition of ECM. They are also the primary target of the only two small-molecule therapeutics with FDA approval to treat IPF, namely the antifibrotics Nintedanib and Pirfenidone(Wollin *et al*., 2015; Margaritopoulos *et al*., 2016; Vancheri *et al*., 2018). Antifibrotics aim to reduce the development of fibrosis, specifically by inhibiting profibrotic signaling and thus restricting the differentiation of fibroblast to the highly active form of fibroblasts responsible for excessive collagen production and expression of α-smooth muscle actin (α-SMA) known as the myofibroblast(Margaritopoulos *et al*., 2016, 2016; Barratt *et al*., 2018; Sgalla *et al*., 2018; Vancheri *et al*., 2018). Despite the approved use of these antifibrotic compounds in 2014, their efficacy is limited, and their side effects are often severe (Noble *et al*., 2011; Vancheri *et al*., 2018; Flaherty *et al*., 2019; Aimo *et al*., 2020; Sgalla *et al*., 2021). Moreover, there are only a tiny handful of compounds in Phase III clinical trials, suggesting gaps in IPF drug delivery and design(Sgalla *et al*., 2021). Specifically, the lack of physiological mimicry by *in vitro* model systems has dramatically reduced our ability to test and produce efficacious FDA-approved therapies(Ahluwalia *et al*., 2014).

Many discovery programs for antifibrotic therapeutics begin with 2-dimensional (2D) tissue cultures *in vitro* models; however, a significant disadvantage of 2D cultures is the absence of the cell-ECM interactions that are known to be pivotal for cell differentiation, gene and protein expression, responsiveness to stimuli, and other critical cellular functions during fibrosis(Kapałczyńska *et al*., 2018; Evangelista-Leite *et al*., 2021; Woodley *et al*., 2021). We have previously demonstrated that fibroblasts extracted from patients with advanced-stage IPF (IPF-F) and normal human lung fibroblasts (NHLF) undergo a dramatic phenotypic change when removed from the complex lung microenvironment and transferred to stiff tissue culture plastic(Emblom-Callahan *et al*., 2010; Rodriguez *et al*., 2018). When cultured on 2D plastic, diseased and non-diseased fibroblasts synchronize into an active, smooth-muscle-actin-expressing phenotype, repressing the expression of disease-associated characteristics. To ensure precise evaluation during the preclinical stage of therapeutic development, it is essential to replicate the disease microenvironment’s pathological features accurately. This involves replicating the mechanical and biochemical signals from the fibrotic extracellular matrix (ECM), which have been demonstrated to promote disease progression^18–20^. Therefore, it is crucial to analyze diseased cells in a microenvironment that accurately mimics these signals for assessment. The ECM is a rich cytokine source strongly influencing fibroblasts’ migration, proliferation, and activity(de Hilster *et al*., 2020; Evangelista-Leite *et al*., 2021; Woodley *et al*., 2021). Recent advancements in *in vitro* disease models are improving our understanding of fibroblast interaction with *in vivo* ECM components, enzymes, and different cell types *in vivo.* However, a knowledge gap persists in how the diseased matrix regulates the behavior of fibroblasts and their communication with other cells in the fibrotic microenvironment(Huang *et al*., 2012a; Herrera *et al*.).

Beyond the fibroblast and their pathological contribution to disease progression, there are also immune cells to consider in this milieu. These cells include monocytes and their tissue-resident counterparts, macrophages, which have been recognized to play a significant role in IPF pathogenesis. Macrophages are one of the main reservoirs of profibrotic cytokines, including transforming growth factor beta (TGFβ), a primary effector molecule in fibrosis and regulator in fibroblast activation(Zhang *et al*., 2018; Cheng *et al*., 2021; Ogawa *et al*., 2021). Many studies have implicated macrophages derived from circulating monocytes, which accumulate in the fibrotic lung microenvironment, in the pathogenesis of IPF(Sica and Mantovani, 2012; Arora *et al*., 2018; Zhang *et al*., 2018; Kishore and Petrek, 2021; Ogawa *et al*., 2021). To fortify the evidence, other studies have indicated that elevated levels of circulating monocytes in the peripheral blood of patients with idiopathic pulmonary fibrosis (IPF) are positively associated with accelerated disease progression and elevated mortality risk. These findings underscore the significant role of monocytes in the pathogenesis of IPF(Scott *et al*., 2019).

To better understand the role of immune cells in IPF pathogenesis, a few recent studies have focused on generating innovative *in vitro* systems in which fibroblasts are co-cultured with macrophages in 3D collagen matrices(Ullm *et al*., 2020; Novak *et al*., 2023). For example, Novak *et al*. demonstrated that direct co-cultures of fibroblasts and macrophages in 3D collagen matrices result in altered crosstalk due to the proximity of the cells, suggesting that collagen matrices are a suitable platform for *in vitro* crosstalk studies(Novak *et al*., 2023). Additionally, Ullm *et al*. found that macrophage and fibroblast co-culture systems using 3D collagen matrices result in a significant secretion of interleukin-10 (IL-10) from macrophages, eliciting fibroblast to myofibroblast differentiation(Ullm *et al*., 2020). IL-10 is an anti-inflammatory cytokine that plays a crucial role in the immune response, and in IPF, it has been implicated as a fibrotic modulator and potential therapeutic target(Millar, 2006; Foster *et al*., 2015; Greiffo *et al*., 2016; She *et al*., 2021). These studies suggest that 3D matrix co-cultures form highly relevant biomimetic models for the late stages of wound healing, enabling the comprehensive investigation of fibroblast-immune crosstalk. These findings indicate the importance of matrix and cell-to-cell signaling, thus opening a new area of research around the ECM and its influence on disease progression.

In the past few decades, *in vitro* models have improved to meet the growing need for model systems that effectively mimic the complex *in vivo* fibrotic microenvironment. For example, 3D collagen hydrogel models with tunable mechanical stiffness have been applied to elucidate fibroblasts’ mechanoregulation(Smithmyer *et al*., 2014; Kapałczyńska *et al*., 2018; Woodley *et al*., 2021; Vazquez-Armendariz *et al*., 2022). Collagen, however, is only one example of a material used to generate a 3D biomimetic system. Currently, models range from naturally derived native ECM components to synthetic tunable materials that aim to resemble the composition of the *in vivo* environment(Smithmyer *et al*., 2014; Matera *et al*., 2020; Woodley *et al*., 2021). Multiple studies have observed that in 3D models, including naturally derived or synthetic hydrogels, fibroblasts transition into a quiescent state that resembles their proliferative activity *in vivo* prior to activation and entry into wound healing(Kono *et al*., 1990; Woodley *et al*., 2021). While these models are a promising tool to investigate *in vivo* signaling, they do not reflect the complexities of diseases such as IPF, which encompasses elements including cells, their environment, their interaction with immune cells, and their matrix. Implementing 3D hydrogels for fibroblast research is an effective and necessary advance to recapitulate disease-relevant conditions *in vitro*(Evangelista-Leite *et al*., 2021).

The advent of polymer-based 3D hydrogels, such as those made from collagen, has allowed the interrogation of understudied disease-relevant properties, most notably fibroblast contractility (Mattyasovszky *et al*., 2017; Denton *et al*., 2018; Kumari *et al*., 2022). These 3D models demonstrate that fibroblast-mediated hydrogel contraction increases in the presence of profibrotic mediators such as TGFβ, and that this contraction can be reduced using therapeutics Nintedanib and Pirfenidone(Kumari *et al*., 2022). In addition to antifibrotic drugs, anti-inflammatory compounds such as interleukin 4 (IL-4) and interleukin 6 (IL-6) inhibitors have been shown to reduce hydrogel contraction(Mattyasovszky *et al*., 2017; Denton *et al*., 2018). Hydrogels demonstrate the potential to provide a suitable platform for investigating the behavior of fibroblasts in a realistic environment. This method can also help elucidate the incompletely understood role of the immune system in the progression of fibrosis. However, it must be noted that this approach is still in its early stages and requires further optimization.

In this work, we present a patient-derived *in vitro* IPF hydrogel model that recapitulates key disease-associated elements necessary to assess the pathobiology of IPF more accurately *in vitro*. First, we quantified the contractility of fibroblasts (both NHLF and IPF-F) when cultured in our hydrogels. We then observed that fibroblast behavior cultured in our 3D IPF ECM hydrogels exhibited significantly decreased activation and proliferation compared to fibroblasts grown in 2D on tissue culture dishes. Finally, we present evidence that the 3D ECM-derived hydrogel culture induces a proinflammatory state in the fibroblasts. This inflammatory state is validated by a gene enrichment analysis of the proinflammation genes “lost” in 2D culture and by data showing increased inflammatory activity of immune cells in fibroblast-conditioned media. These results suggest a potential application of our hydrogel platform for interrogating fibroblast-macrophage crosstalk and its role in IPF pathogenesis. By characterizing how fibroblasts behave in our patient-derived model of the fibrotic lung, we aim to expand knowledge and provide novel *in vitro* tools for developing more efficacious therapies to treat IPF.

## 2. METHODS

### 2.1 Donor consent and internal review board approval

This study was conducted in accordance with principles set forth by the Declaration of Helsinki. IPF lung tissue was obtained through Inova Fairfax Hospital (Fairfax, VA). All normal control lungs were obtained through the Washington Regional Transplant Community (WRTC). Appropriate written informed consent was obtained from each patient and donor by Inova Fairfax Hospital and the WRTC. This study was approved by the Inova Fairfax Hospital Internal Review Board (IRB #06.083) and the George Mason University Human Subject Review Board (Exemption #5022). All experiments were performed in accordance with relevant guidelines and regulations.

**Table 1:**
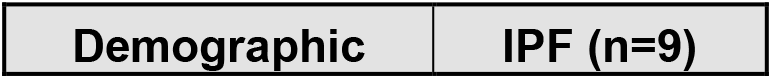
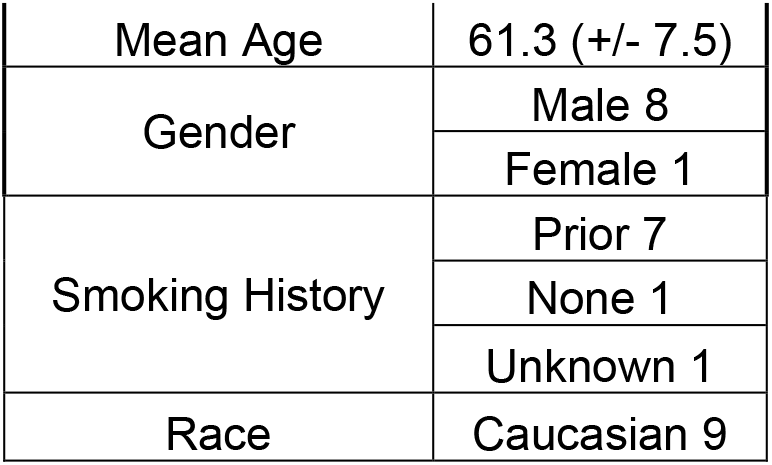
Demographics of all donors of IPF fibroblasts and lung tissue used in this study.

| Demographic | IPF (n=9) |
| --- | --- |
| Mean Age | 61.3 (+/- 7.5) |
| Gender | Male 8 |
|  | Female 1 |
| Smoking History | Prior 7 |
|  | None 1 |
|  | Unknown 1 |
| Race | Caucasian 9 |

### 2.2 Tissue Decellularization

This protocol is adapted from a previously published protocol (Freytes *et al*., 2008) and summarized in Figure 1. IPF lung tissue (n=4) was minced into 2-5 mm^3^ pieces and then washed with room-temperature 1% aqueous solutions of sodium dodecyl sulfate (SDS) and Triton X-100 for 24 h, replacing each solution three times. The tissue was then rinsed twice with 1X phosphate-buffered saline (PBS) and washed with the following solutions for 30 min with two rinses of PBS and seven rinses of water in between: 10 U/mL of DNase in 1X PBS at 37°C, 1 M NaCl for 1 h, and 70% ethanol finalized with all rinses. The scaffold was maintained in PBS solution at 4°C until lyophilization.

**Figure 1.**
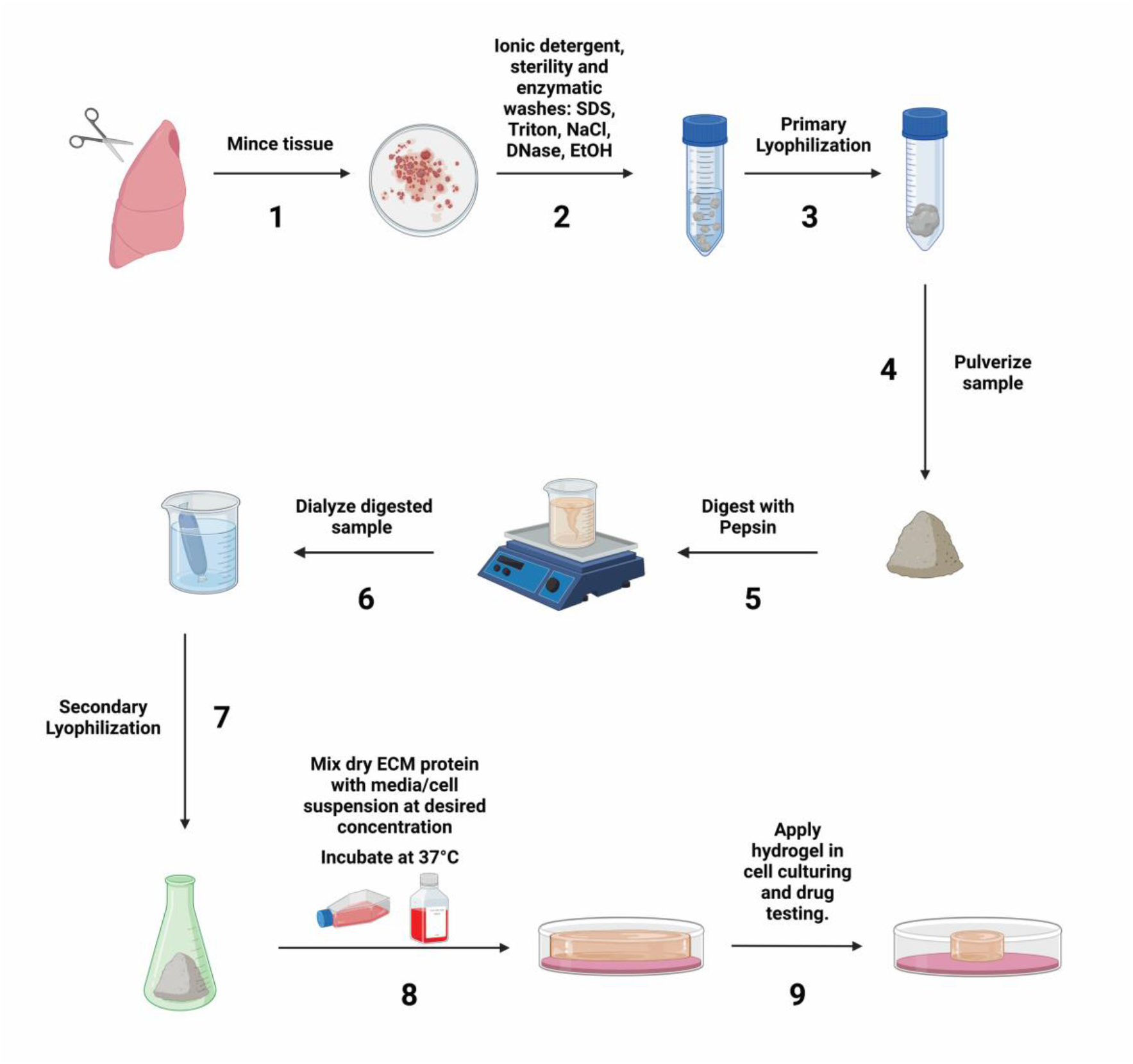
Schematic of IPF ECM hydrogel synthesis protocol. Procured human lung tissue is minced (1) and decellularized with ionic detergent washes and DNase to remove extracellular DNA (2). Scaffold is sterilized with ethanol followed by a lyophilization (3). The scaffold is pulverized (4) and then digested with pepsin (5), followed by a dialysis (6), and finalized by a secondary lyophilization (7). Powder is resuspended with culture media and seeded with cells. Gelation of sample occurs at 37°C (8,9).

**Figure 2.**
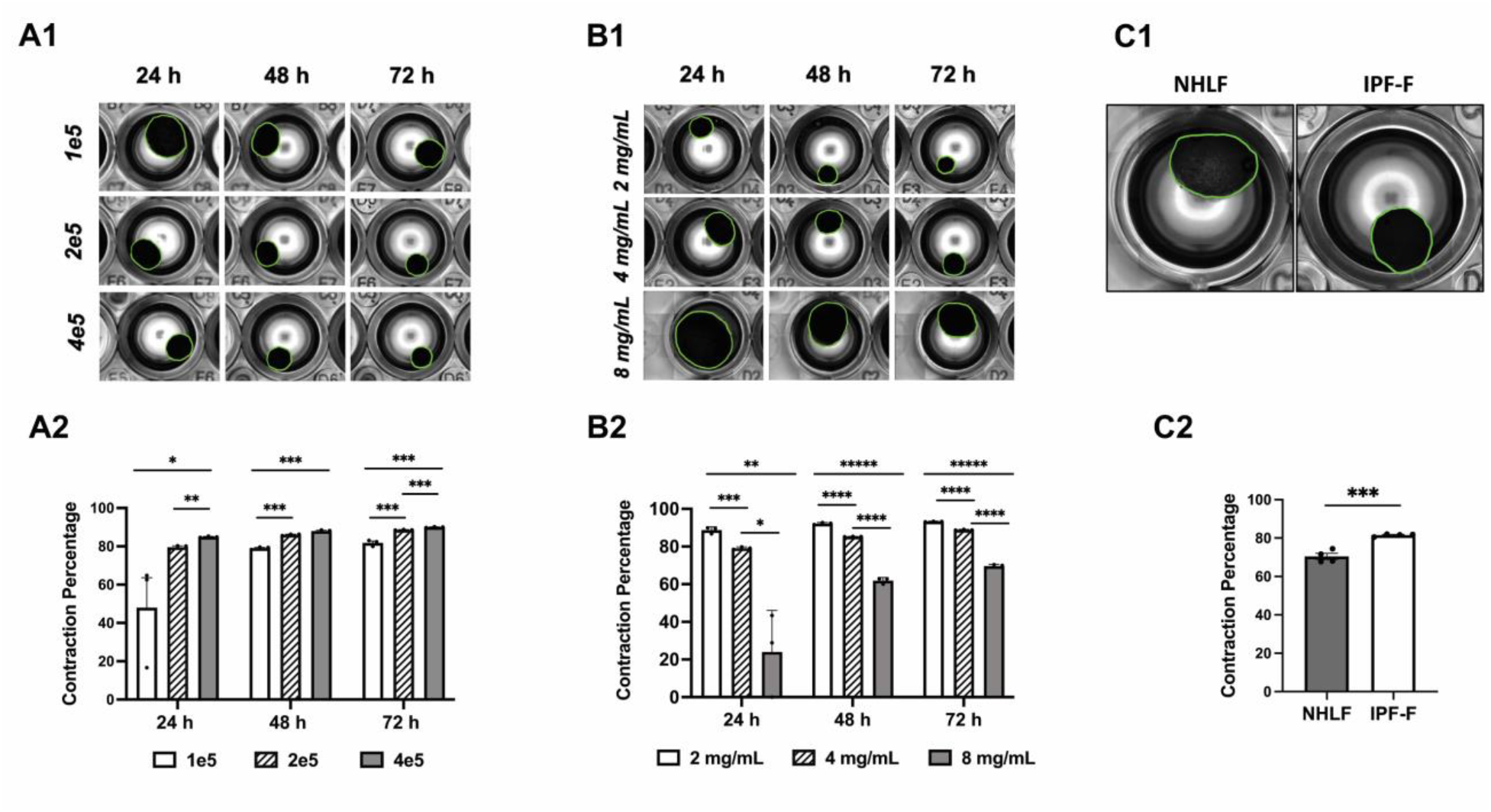
Contraction Rate of IPF Derived Hydrogels. (A1) IPF hydrogel (4 mg/mL ECM protein concentration) seeded with varying fibroblast densities: 1e5, 2e5, and 4e5 cells/well. (A2) Bar graph demonstrating a significant increase at all time points in hydrogel contraction as cell density increases. (B1) IPF hydrogel at varying ECM protein concentrations: 2 mg/mL, 4 mg/mL, and 8 mg/mL with a fibroblast density of 2e5 cells/well. (B2) Bar graph demonstrating a significant decrease of hydrogel contractions at all time points as concentration of IPF ECM in the hydrogel increases. (C1) IPF hydrogel seeded with NHLF and IPF-F at a density of 2e5 and imaged after 24 h. (C2) Significantly increased contraction of IPF-F in comparison to NHLF in IPF hydrogels after 24 h. *: p<0.05, **: p<0.005, ***: p<0.0005, ****: p<0.00005, *****: p<0.000005.

### 2.3 Synthesis of ECM hydrogel

IPF lung scaffold was frozen overnight at –80°C prior to lyophilization. The frozen scaffold was then lyophilized overnight in a Millrock Technology Benchtop Manifold Freeze Dryer. Post-lyophilization, the sample was pulverized with a Spex SamplePrep Freezer/Mill 6875. The powdered lung scaffold was then digested by mixing 200 mg of scaffold powder and 100 mg of pepsin (MP Bio, 02102598-CF) (2:1 ratio) in 20 mL of hydrochloric acid (HCl) at pH 2.0 for 72 h. The pH of the solution was measured at the beginning of the digestion and once every 24 h for a period of 72 h; it was adjusted to pH 2.0 by droplet addition of 12 M HCl. The ECM solution was then neutralized to pH 7.0 with 1 M NaOH, dialyzed with water, changed every hour for a total of 3 times, and left dialyzing overnight. The solution was transferred to a conical tube and frozen at –80°C overnight, followed by a lyophilization for 24 h. The final product, which has the consistency of a spongelike material, was resuspended in a cell culture medium of choice at the desired concentration to form a pre-gel solution. Pre-gel solution is a viscous fluid that is pipetted into a well dish and incubated at 37°C for at least 45 min before reaching gelation. A graphical depiction of the process is presented in Figure 1.

### 2.4 Specimen procurement/dissection and cell culture

The primary fibroblasts (NHLF and IPF-F) used in this study were isolated from human lungs procured in the operating room within minutes of explant. The lungs were oriented from apex to base, and all lung samples used in this study for fibroblast isolation were taken from the peripheral lower lobe of the lung. Fibroblasts were isolated from the lung tissue of patients with advanced-stage IPF (IPF-F) and normal healthy lungs (NHLF) using differential binding as previously described(Emblom-Callahan *et al*., 2010). IPF-F (n=5) and NHLF (n=4) were used in this study. Fibroblasts were cultured in a T-75 flask in Dulbecco’s Modified Eagle Medium (DMEM) (Cytiva, SH30081.05) with 5% fetal bovine serum (FBS) (Atlanta Biologicals, D18043) and 1% penicillin/streptomycin, penicillin (100 I.U/mL) and streptomycin (100 MCH/mL) (Corning, 30-002-CI). Cells were cultured at 37°C using 5% CO_2_. Passages of fibroblasts used for this analysis ranged from 4-7. U937 monocytes (ATCC, CRL-1593.2) were cultured in a T-75 flask using RPMI (ThermoFisher, 11875093) 1640 medium with 10% FBS and 1% penicillin/streptomycin and maintained in a humid incubator at 37°C using 5% CO_2_. U937 cells were differentiated to macrophages through a 48-h incubation with 100 ng/mL of phorbol myristate acetate (PMA) (MP Bio, 02151864-CF).

### 2.5 IPF hydrogel cultures and conditioned media experiments

Fibroblasts were split/passaged when cells reached ∼80% confluency using 0.05% trypsin/EDTA (ThermoFisher, 25300054) and pelleted at the desired cell density. Fibroblasts were seeded in a 6-well dish with 2 mL of DMEM complete medium for regular tissue culture and cultured in the incubator for two days. For IPF hydrogel culture, fibroblasts were pelleted at 4 × 10^5^ cells per gel and slowly resuspended in the 600 µL pre-gel solution. The pre-gel with the cell suspension was then pipetted at a volume of 300 µL into a 48-well tissue culture dish and placed in a humidified incubator at 37°C with 5% CO_2_ for 30 minutes to allow gelation before adding 500 µL of DMEM complete medium warmed to 37°C and placing the culture dish in the incubator.

For experiments exposing immune cells to fibroblast conditioned media, the conditioned media was collected and centrifuged at 1000 rpm for 5 min to remove any cell debris and/or floating cells. Conditioned media was used to challenge 5 × 10^5^ U937 cells per gel on a 24-well tissue culture dish for 48 h. For PMA treatments, 2.5 × 10^5^ U937 cells were seeded with 100 ng/mL PMA-supplemented RMPI media and cultured for 48 h followed by RNA isolation. PMA media was removed, and adherent cells were washed with PBS, followed by treatment with fibroblast conditioned media for 48 h and followed by RNA isolation.

### 2.6 RNA isolation and cDNA synthesis

RNA was isolated from IPF-F, NHLF, and U937 cells using the Omega Biotek E.Z.N.A Total RNA Kit I (Omega Biotek, R6834-02). For 3D cell extractions, hydrogel was incubated in complete media with 1% collagenase type I (ThermoFisher, 17100017) for 25-30 minutes. Once the cells disassociated from the hydrogel, the solution was vigorously pipetted until hydrogel clumps were no longer visible. The solution was then centrifuged for 5 min at 1000 rpm, and the pellet was resuspended in lysis buffer followed by isolation according to the manufacturer’s instructions. For analysis, 0.1-0.2 µg of RNA was converted to cDNA using the VERSO cDNA Synthesis Kit (ThermoFisher, AB1453A) prior to quantitative PCR.

### 2.7 Realtime Quantitative PCR

Quantitative PCR (qPCR) was performed in a Bio-Rad CFX Connect Optics Module by using the Applied Biosystems Fast SYBR Green Master Mix kit (ThermoFisher, 4385612). qPCR was normalized to 18S expression using the delta-delta Ct method(Pfaffl, 2001). Statistical analysis for Figure 3 was performed via paired T-test in Microsoft Excel. Welch’s T-test was performed via GraphPad Prism 9 for Figure 4. One-way ANOVA was performed via GraphPad Prism 9 for Figures 5 and 6. Expression fold differences with p-values less than 0.05 were considered significant. A full list of primer sequences is included in Supplemental Table 1.

**Figure 3.**
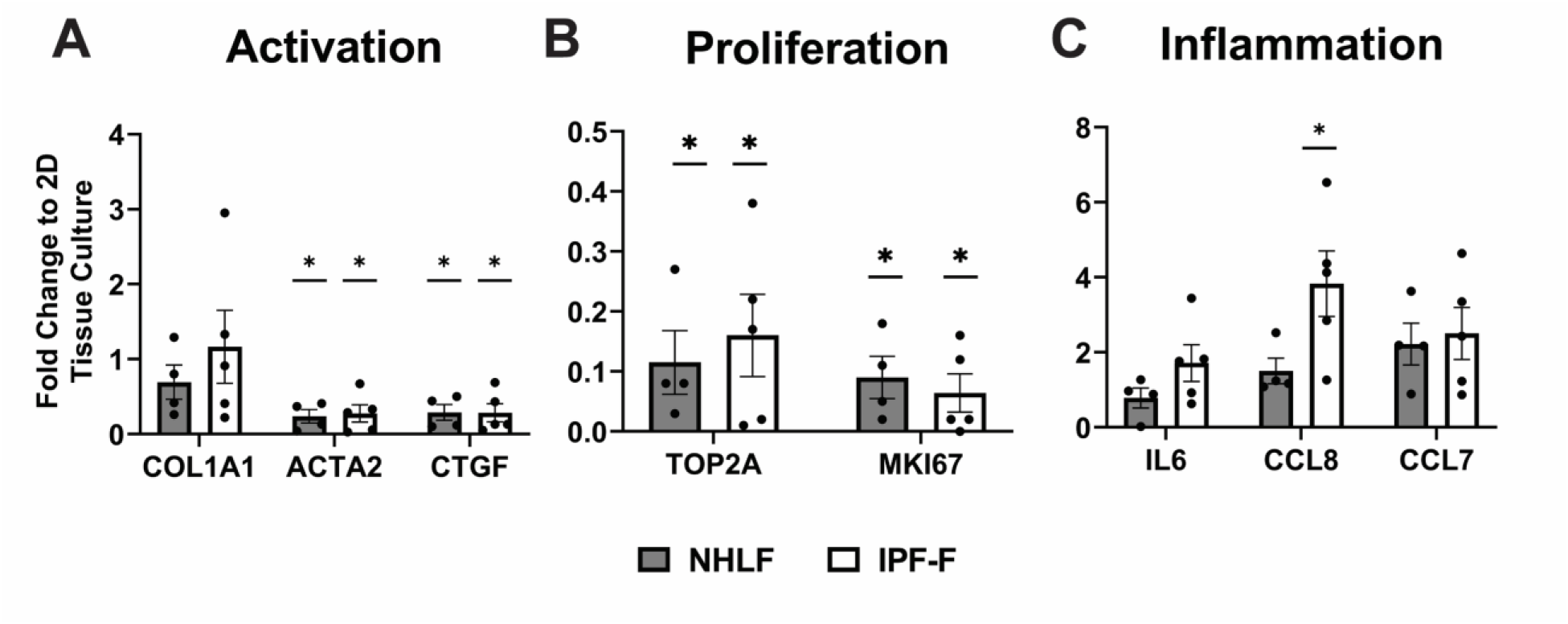
Gene Expression of IPF-F and NHLF Fibroblasts Seeded in IPF Hydrogel and Tissue Culture Plastic: (A) Gene expression of fibroblast activation markers: *Col1a1, ACTA2,* and *CTGF*. (B) Gene expression of ligands associated with inflammation: *IL6, CCL8, and CCL7 (*C) Gene expression of markers associated with cell proliferation: *TOP2A* and *MKI67*. *: p<0.05.

**Figure 4.**
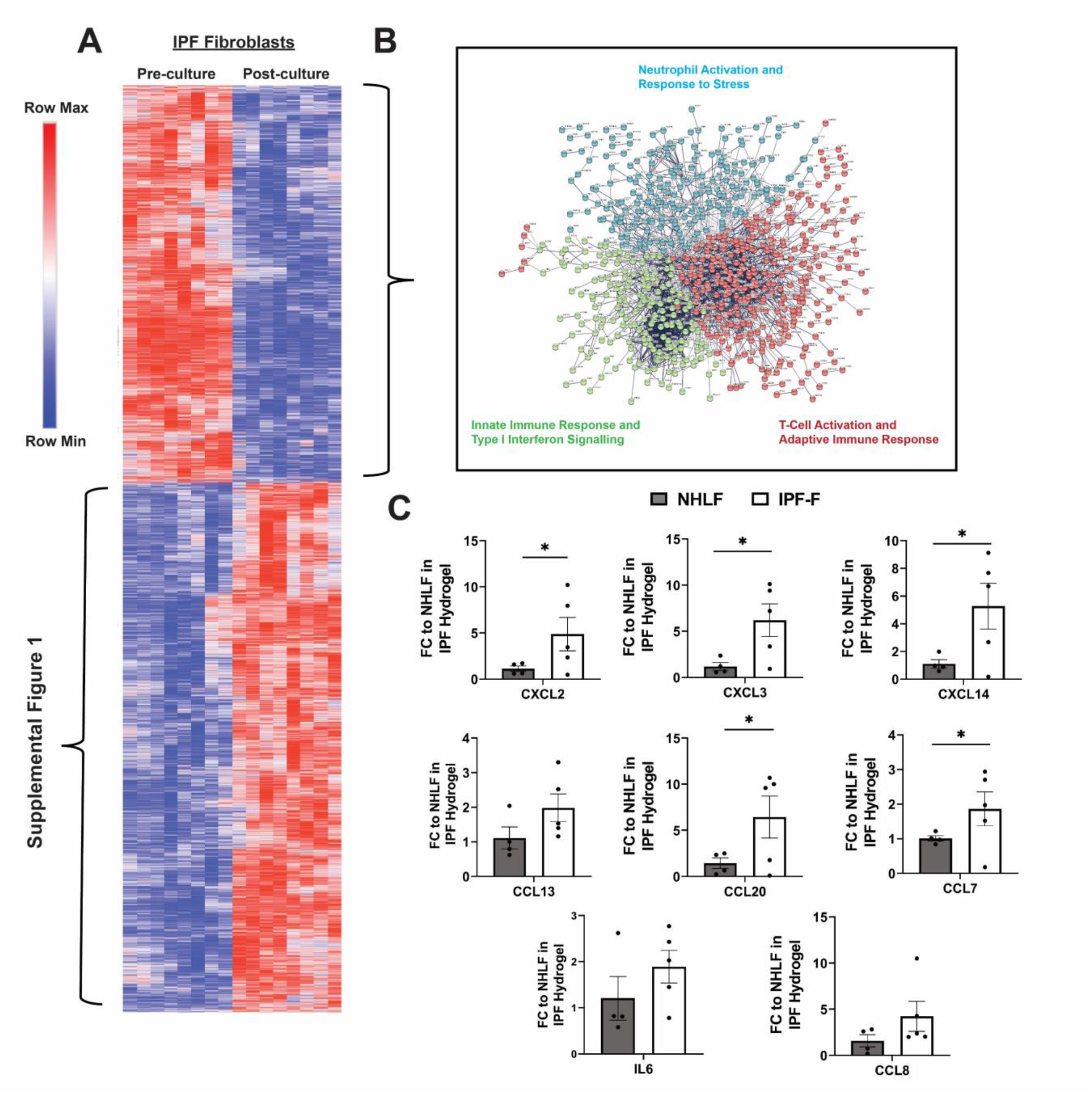
Proinflammatory Gene Expression of Fibroblasts in Fibrotic ECM Hydrogel. (A) Microarray data highlighting gene expression changes of IPF fibroblasts immediately prior to culture and after 3 weeks. (B) STRING analysis of protein interactions compiled from genes in the bottom block of the heat map in Figure 4A. (C) Bar graphs demonstrating fold change of gene expression *(CCL20, CXCL2, CXCL3, CXCL13, CXCL14, CXCR4)* from STRING diagram in Figure 4B comparing NHLF and IPF-F seeded in IPF hydrogels against the average of NHLF expression*. IL6, CCL7* and *CCL8* are not part of the genes in STRING diagram but were identified as inflammatory genes in Figure 3B. *: p<0.05.

**Figure 5.**
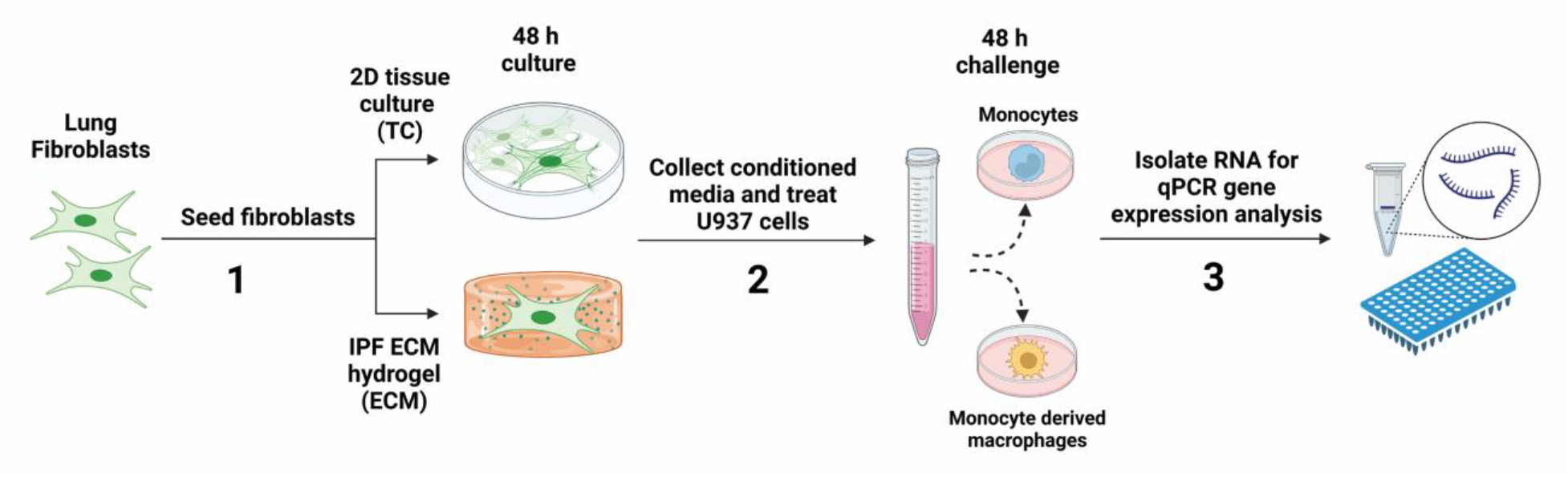
Schematic Depiction of Conditioned Media Experiments. Fibroblasts are seeded in IPF hydrogels for 48 h (1) and conditioned media is collected. and used to challenge U937 (monocytes) and PMA treated U937 (monocyte derived macrophages) (2). After 48 h RNA is isolated (3).

**Figure 6.**
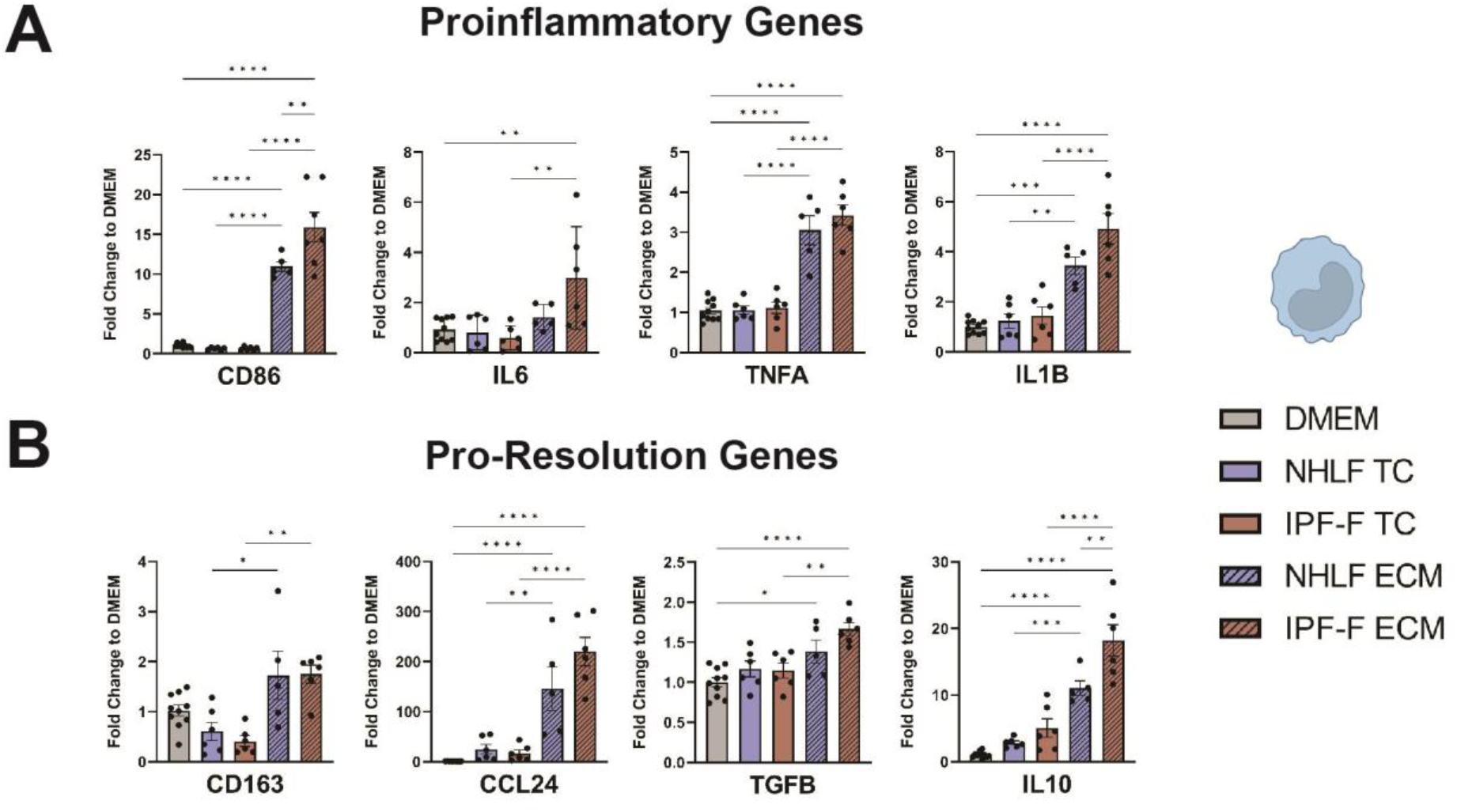
Gene Expression in Monocytes After Fibroblast Conditioned Media Challenge. (A) Gene expression fold change of proinflammatory genes: *CD86*, *IL6*, *TNFA*, and *IL1B* (B) Gene expression fold change of pro-resolution genes: *CD163*, *TGFB*, *CCL24*, *IL10*. (C) Gene expression fold change of markers cell cycle and proliferation: *TOP2A*, *PCNA*, and *P16.* *: p<0.05, **: p<0.005, ***: p<0.0005, ****: p<0.00005.

### 2.8 Immunoblot

Cultured cell samples were lysed in protein lysis buffer (RIPA, protease inhibitor, and phosphatase inhibitors). Protein concentration of samples was determined by BCA assay. 20 µg of each sample were subjected to Sodium dodecyl-sulfate polyacrylamide gel electrophoresis (SDS/PAGE) using precast 10% Bis-Tris NuPage Gels (Invitrogen), transferred onto PVDF membranes, probed overnight with primary antibodies, and finally labeled for 1 hour with a species specific HRP conjugated secondary antibody. Detection was performed using a chemiluminescence substrate (ThermoFisher) and images were acquired using a Bio-Rad Chemidoc imaging system. Primary antibodies against the following proteins were used for immunoblotting: PCNA (Cell signaling) and β-Actin (Cell signaling). Secondary antibodies were goat anti-mouse or rabbit HRP (Cell signaling).

### 2.9 Microscopy and Hydrogel Contraction Analysis

Images of hydrogels were captured with a Life Technologies EVOS FL Auto imaging system in the 48-well vessel layout taken at 4X magnification. The contractility of hydrogels was analyzed by measuring the reduction in surface area of the contracted hydrogel touching the bottom of the well in ImageJ using the oval tool or by freehand outlining of the gel shapes to measure the area of the entire well and that of the hydrogel. Statistical analysis for contractions was performed using one-way paired T-tests in Microsoft Excel. Expression fold differences with p-values less than 0.05 were considered significant. The following equation was used to estimate the percent contraction of the hydrogel surface:

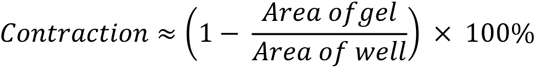

### 2.10 Microarray Analysis

Gene expression of fibroblast transition through culture is a subset of data from previously published work available through GEO accession GSE87175(Rodriguez *et al*., 2018). Protein-protein interactions were obtained using the STRING (Search Tool for Retrieval of Interacting Genes/Proteins) database. Heat map was generated using MORPHEUS from the Broad Institute with application of K means clustering.

### 2.11 Cell Viability Assay

U937 Monocytes were seeded in 96-well plates at a density of 3 × 10^4^ cells/well in 100 mL/well and incubated for 48 h with fibroblast-conditioned media (DMEM). Non-conditioned DMEM was used as a control. CCK-8 (Enzo Life Sciences) was added (10 uL/well) and incubated for 1 h according to the manufacturer’s instructions. The absorbance was measured at 450 nm with a microplate reader (Biotek Cytation 5).

## 3. RESULTS

### 3.1 Hydrogel Contraction Depends on Fibroblast Density, Fibroblast Disease Status, and ECM Protein Concentration

Both IPF-F and NHLF exhibited contractility when cultured in our IPF-patient-derived hydrogels. The extent of hydrogel contraction was estimated as an *in vitro* proxy for fibroblast contractility *in vivo*. We conducted various experiments to quantify the differential effects of fibroblast density and ECM protein concentration on hydrogel contraction.

To evaluate the effect of cell seeding density on contraction, we seeded fibroblasts in our ECM-derived hydrogels at concentrations of 1 × 10^5^ (1e5), 2 × 10^5^ (2e5), and 4 × 10^5^(4e5) cells per hydrogel (ECM protein concentration 4 mg/mL in each case). We initially observed contraction 24 h after seeding and detected a significant difference among all three densities by 72 h (Figure 2A). For all cell densities, the contraction percentage plateaus at just over 80% (i.e., the hydrogels shrink to less than 20% of their original size); 80% contraction is reached just after 24 h in gels seeded with 2e5 and 4e5 fibroblasts, but only after 72 h with 1e5 fibroblasts.

To evaluate the impact of ECM protein concentration on contraction, we seeded 2e5 fibroblasts in hydrogels with ECM protein concentrations of 2, 4, and 8 mg/mL. Within 24 h, contraction was observed in 2 and 4 mg/mL hydrogels, with a significant difference between 4 mg/mL and 8 mg/mL at 48 h (Figure 2B).

Finally, we compared the contractile capacity of NHLF and IPF-F in 4 mg/mL hydrogels (2e5 cells per hydrogel). We observed significantly higher contraction in IPF-F-seeded hydrogels after 24 h (Figure 2C). The fibroblast cell density of 2e5 seeded in 4 mg/mL ECM protein concentration yielded the best consistency in contraction profiling. Given these results, all further experiments employed a cell density of 2e5 and an ECM protein concentration of 4 mg/mL.

### 3.2 Fibroblasts Seeded in ECM Hydrogels Reduce Activation and Proliferative Gene Expression

To evaluate the fibroblast phenotype in 3D culture, we performed qPCR using markers associated with fibroblast activation during wound healing, inflammatory signaling, and proliferation. By analyzing genes associated with fibroblast activation, including Collagen 1a1 (*COL1A1),* Smooth-Muscle-Actin (*ACTA2),* and Connective Tissue Growth Factor *(CTGF)*, we identified a significant decrease in the activation signature of fibroblasts seeded in IPF hydrogels as compared to standard tissue culture plastic (Figure 3A). These reductions in gene expression can be attributed to decreased mechanical stiffness of the IPF-derived hydrogel compared to tissue culture plastic, the latter of which is a well-characterized driver of the active myofibroblast phenotype(Dasgupta and McCollum, 2019; Scott *et al*., 2021). Further supporting this observation, we also observed a significant decrease in proliferating cell nuclear antigen (*PCNA*) and type IIA topoisomerase (*TOP2A*), genes associated with proliferation in both IPF-F and NHLF (Figure 3B). We also looked at the expression of the inflammation-associated genes Interleukin 6 (*IL6*), Chemokine C-C motif ligand 8 (*CCL8*), and Chemokine C-C motif ligand 7 (*CCL7*)(Burgess *et al*., 2016; Hamai *et al*., 2016) to assess the possible proinflammatory effects of seeding cells in the IPF hydrogels (Figure 3C). While only *CCL8* had a significantly increased expression in IPF-F, this initial screening of inflammation genes presented an intriguing avenue for further characterization.

### 3.3 IPF Fibroblasts Seeded in Fibrotic ECM Enter a Proinflammatory State

We have previously reported that gene expression of freshly isolated lung fibroblasts is dramatically altered through extended tissue culture. These changes in NHLF and IPF-F result in a convergent phenotype marked by high activation compared to the *in-vivo* state(Rodriguez *et al*., 2018). In our previous work, we identified a subset of genes that were altered in both IPF-F and NHLF as the fibroblasts transitioned to tissue culture plastic. We attributed those alterations to 2 factors: 1) the absence of biochemical cues found *in vivo* and 2) the stiffness of tissue culture plastic. A heat map of those genes highlights two subsets of genes altered through tissue culture, with one set increasing and one set decreasing (Figure 4A). Evaluation of genes upregulated upon transfer to tissue culture plastic is described in Supplemental Figure S1; however, our focus in this manuscript is on the genes that were downregulated or lost upon transition to traditional 2D tissue culture (roughly the top half of Figure 4A). Thus, we performed gene enrichment analysis on these downregulated genes using StringDB to identify key pathways not described in the previous work. Crucially, we identified a major hub of genes associated with inflammatory signaling that are downregulated in tissue culture plastic (Figure 4B).

Paired with our initial observations suggesting a potential increase of proinflammatory signaling for cells seeded in fibrotic ECM (Figure 3C), we hypothesized that the fibrotic ECM may “rescue” the inflammatory signaling lost upon transition to 2D tissue culture. We chose six genes from the STRING analysis: Chemokine (C-C motif) ligand 20 (*CCL20*), Chemokine (C-X-C motif) ligand 2 *(CXCL2),* Chemokine (C-X-C motif) ligand 3 (*CXCL3),* Chemokine (C-C motif) ligand 13 (*CCL13),* Chemokine (C-X-C motif) ligand 14 (*CXCL14),* and the three other inflammatory genes from the Figure 3C: *IL6, CCL7* and *CCL8*. Comparing NHLF and IPF-F seeded in our hydrogels, we observed that IPF-F exhibited significantly more gene expression for many of the selected cytokines (Figure 4C). This data further supports a potential proinflammatory nature of the fibrotic ECM and suggests that IPF-F have an increased sensitivity to biochemical cues within our IPF patient-derived ECM hydrogel matrices, compared to 2D tissue culture plastic.

### 3.4 Conditioned Media from Fibroblasts in Fibrotic ECM Activate Monocytes and Macrophages

To further examine the proinflammatory state of the fibroblasts in IPF ECM, we developed a conditioned media experiment using the monocyte cell line U937 and monocyte-derived macrophages (U937s treated for 48 h with 100 ng/mL PMA). We hypothesized that the increased expression of inflammatory cytokines observed in the fibrotic hydrogels would translate to increased release of cytokines to the surrounding media of the fibroblast. Furthermore, we expected this cytokine-enriched media to elicit a potent response in either monocytes or macrophages. The assay protocol is thoroughly described in our Methods section and graphically depicted in Figure 5.

With our primary interest being fibrosis, we broadly defined cytokine panels that have previously been associated with either the inflammatory (*CD86, IL6, TNFA* and *IL1B)* or resolution (*CD163, TGFB, CCL24, and IL10)* phenotypes of macrophages in the fibrotic lung(Arora *et al*., 2018; Zhang *et al*., 2018; Cheng *et al*., 2021). Additionally, we investigated whether cytokines released from fibroblasts could promote the expression of a proliferative gene set in either monocytes or macrophages. Conditioned media from fibroblasts seeded in traditional 2D tissue culture had no significant effect on monocyte gene expression across all genes tested (Figure 6A-C). However, conditioned media from NHLF and IPF-F cultured in IPF hydrogels induced significant increases in proinflammatory and proresolution gene expression in monocytes (Figure 6A-B). The expression of select genes, CD86 and IL10, were significantly higher in monocytes treated with IPF-F conditioned media than in NHLF-conditioned media, further supporting our initial observation of an increased proinflammatory profile (Figure 4). This also suggests that IPF-F may enter a more inflammatory state in fibrotic ECM than NHLF.

Experiments using macrophages in co-culture demonstrate a differential effect when comparing conditioned media from 2D tissue culture (TC) and 3D ECM cultured fibroblasts. The proinflammatory signature was again increased in macrophages treated with ECM conditioned media compared to both DMEM and TC conditioned media (Figure 7A). Surprisingly, apart from *CCL24*, which was significantly upregulated by ECM conditioned media, this increased expression was not observed for the profibrotic gene signature. In the case of *IL10* and *CD163*, both conditioned media permutations resulted in a decrease in the expression of these genes, with no significant difference in *TGFB* (Figure 7B).

**Figure 7:**
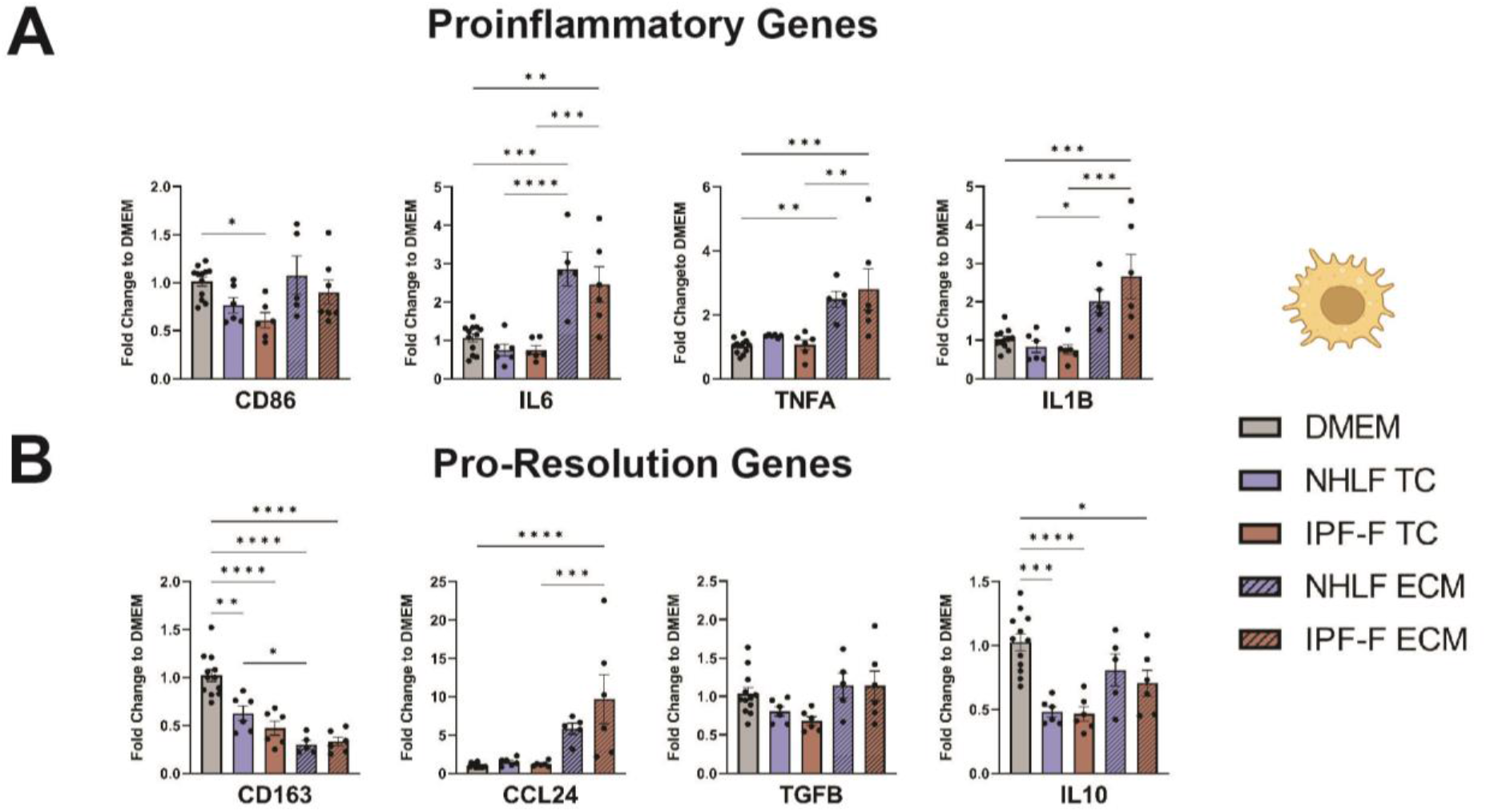
Gene Expression in Monocyte-Derived Macrophages After Fibroblast Conditioned Media Challenge. (A) Gene expression fold change of proinflammatory genes: *CD86*, *IL6*, *TNFA*, and *IL1B* (B) Gene expression fold change of markers cell cycle and proliferation: *TOP2A*, *PCNA*, and *P16.* (C) Gene expression fold change of pro-resolution genes: *CD163*, *TGFB*, *CCL24*, *IL10*. *: p<0.05, **: p<0.005, ***: p<0.0005, ****: p<0.00005.

### 3.5 Conditioned Media from Fibroblasts in Fibrotic ECM Increases Immune Cell Proliferation

Recognizing the known role of monocytes in IPF and their correlation with disease progression(Scott *et al*., 2019; Kreuter *et al*., 2021), we investigated whether our system altered the immune cells’ proliferative state. We found that conditioned media from IPF-F, but not NHLF, induces the expression of a proliferative gene signature in monocytes (Figure 8A). In contrast to the effect on monocytes, ECM-derived conditioned media does not induce the expression of proliferation-associated genes in macrophages (Figure 8B). Immunoblot analysis of monocytes exposed to fibroblast conditioned media confirms increased PCNA in monocytes exposed to conditioned media derived from all ECM conditions. Finally, functional measurement of proliferation using the CCK8 assay confirms that ECM derived fibroblast conditioned media significantly increases monocyte proliferation (Figure 8E).

**Figure 8:**
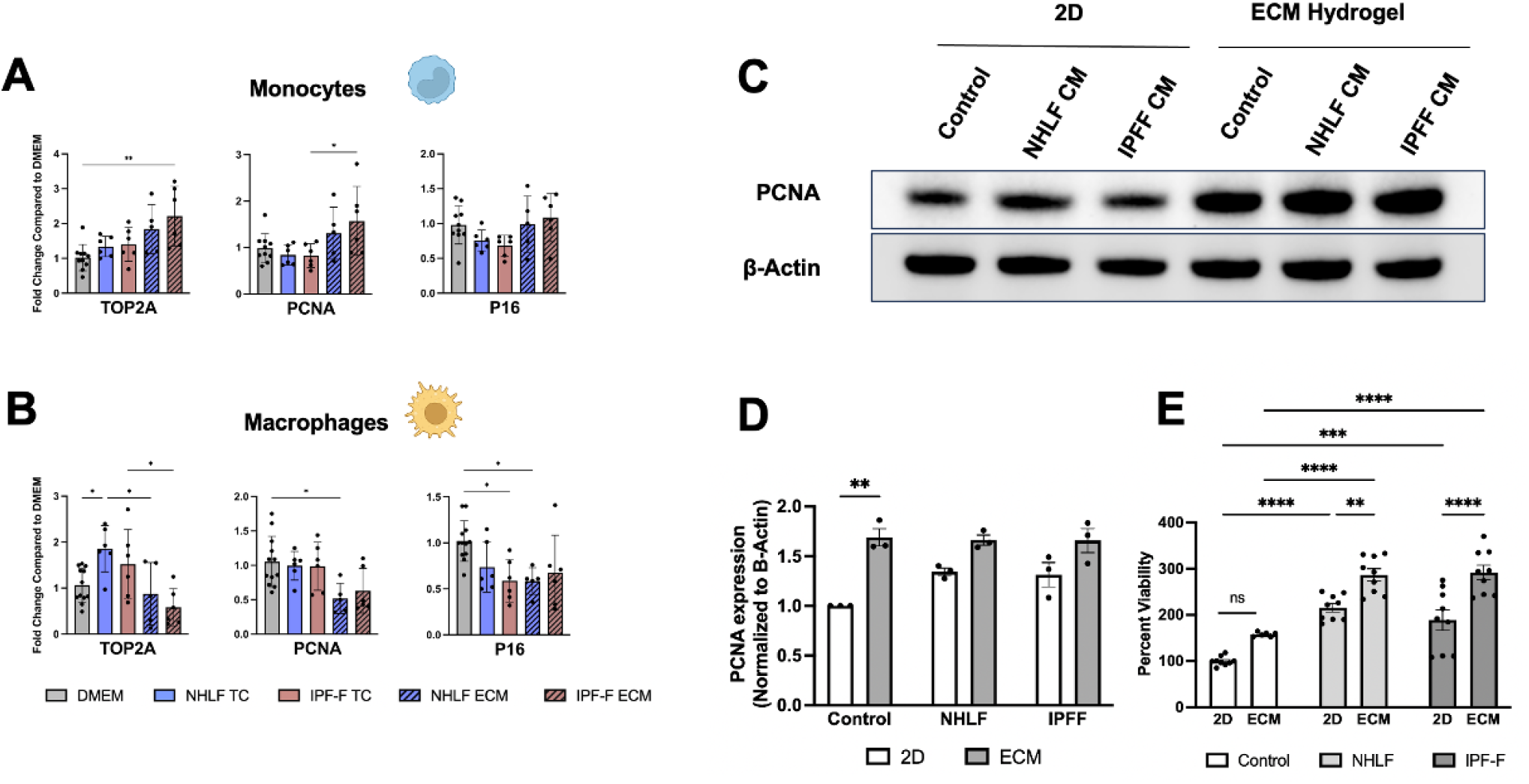
Proliferation of Monocytes After 48 h Fibroblast Conditioned Media Challenge. *(A)* Gene expression fold change of markers of cell cycle and proliferation in U937 monocytes: *TOP2A*, *PCNA*, and *P16.* (B) Gene expression fold change of markers cell cycle and proliferation in U937 monocyte-derived macrophages: *TOP2A*, *PCNA*, and *P16. (C)* Immunoblotting image of PCNA and β-Actin on U937 lysates exposed to fibroblast conditioned media for 48 h. (D) PCNA protein expression normalized to β-Actin of immunoblot in Figure 8C is shown as a bar graph. (E) Percent viability of U937 monocyte exposed to fibroblast conditioned media (CCK8 assay) shown as a bar graph; non-conditioned medium was used as a control. *: p<0.05, **: p<0.005, ***: p<0.0005, ****: p<0.00005.

**Figure 9:**
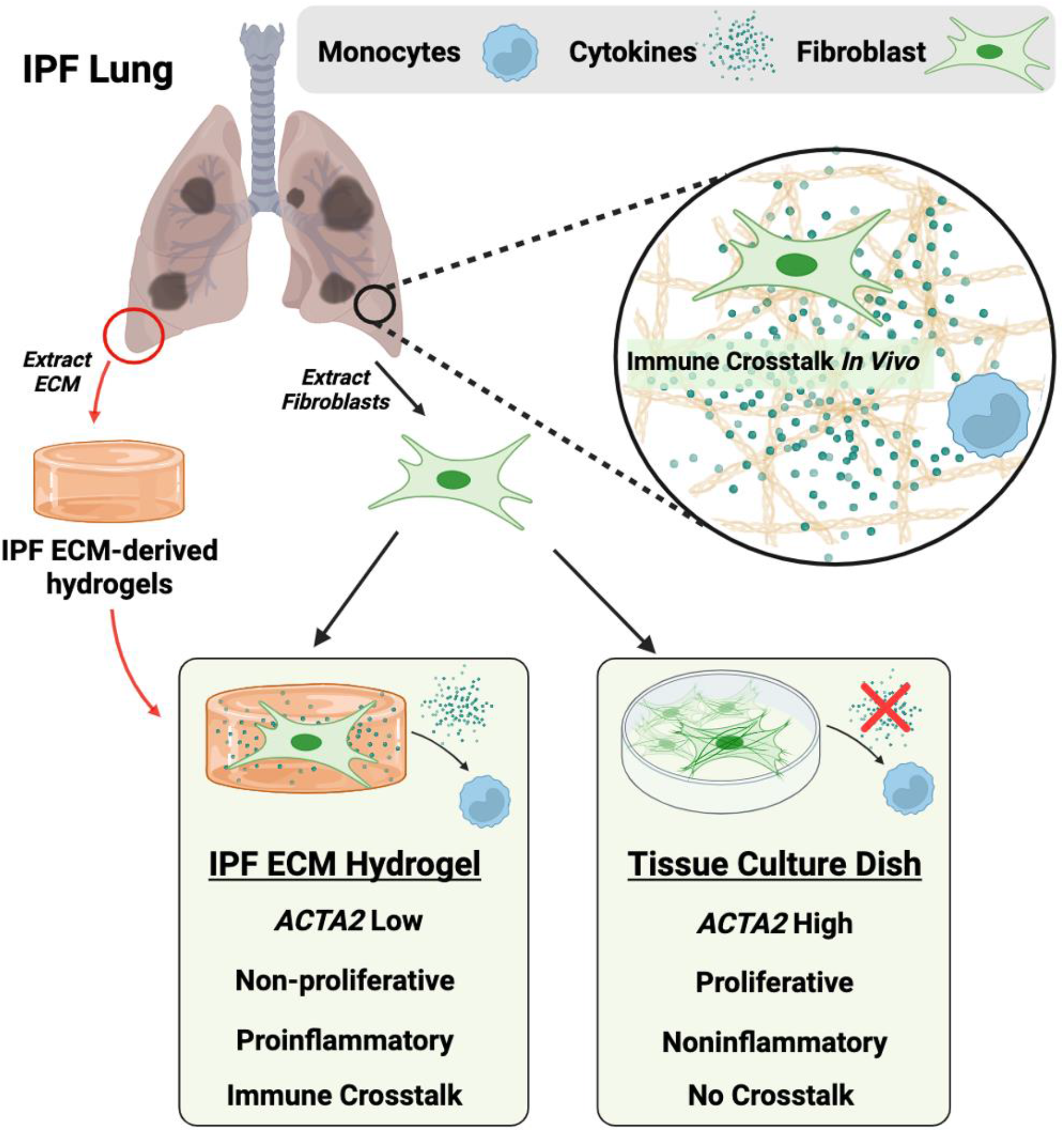
Immune-fibroblast crosstalk in the IPF lung is a poorly understood aspect of disease biology that is difficult to model *in-vitro*. Fibroblasts isolated from the IPF lungs and seeded into IPF ECM hydrogels maintain high levels of cytokine expression as compared to fibroblasts seeded on tissue culture plastic. These cytokines are potent activators of both monocytes and monocyte-derived macrophages.

## 4. DISCUSSION

Since the first studies demonstrating the differential response of fibroblasts to alterations in surface stiffness(Arora *et al*., 1999), our growing understanding of the mechanosensing nature of the fibroblast has been facilitated by applying tunable matrices of varying stiffness(Wipff and Hinz, 2009; Huang *et al*., 2012b; Liu *et al*., 2014; Smithmyer *et al*., 2014; Matera *et al*., 2020). The overwhelming observation, documented in numerous studies over more than 20 years, has been that stiff matrices induce a myofibroblast state marked by α-SMA expression, and this phenotype is lost when fibroblasts are returned to a soft matrix(Dasgupta and McCollum, 2019; D’Urso and Kurniawan, 2020; Matera *et al*., 2020; Scott *et al*., 2021). Recent advances in single-cell technology have uncovered that lung mesenchyme is not a heterogeneous population during wound healing or fibrosis stages. Fibroblast populations in the lung are now known to be variable, with their local microenvironments playing a crucial role in defining their characteristics rather than the expression of α-SMA(Zepp *et al*., 2017; Adams *et al*., 2019; Reyfman *et al*., 2019; Tsukui *et al*., 2020; Buechler *et al*., 2021). Our research utilizing ECM hydrogels has revealed that fibroblasts cultured in this environment demonstrate a less proliferative and less α-SMA-expressing phenotype, but we have also observed a significant increase in inflammatory behavior, which suggests that our hydrogel system may help investigate cytokine secretory fibroblasts populations that drive the disease, *in vitro*. By applying hydrogels made from patient-derived fibrotic lung tissue, we contribute to the field in two important ways. First, we establish a unique fibroblast phenotype characterized by decreased proliferative activity and decreased α-SMA gene expression. Second, we report that the inclusion of IPF ECM proteins derived from the hydrogel can alter the behavior of fibroblasts, specifically by promoting inflammatory signatures in NHLF and IPF-F that were not previously observable *in vitro* (Figure 3B and Figure 4C) but known to be essential for IPF pathogenesis *in vivo*. Together, these observations highlight the necessity for developing *in vitro* models with tunable stiffness and a biologically relevant matrix composition.

Through our characterization of this system, we observed that the contractile capacity of fibroblasts depends on two factors: the number of cells in the hydrogel and the concentration of the matrix protein (Figure 2). These observations suggest that increased ECM protein concentration in fibrotic gels increases the hindrance of the hydrogel’s contractile forces exerted by fibroblast and that the hydrogel contractions can be increased by seeding more cells. We also observed that IPF-F have a greater capacity to contract the matrix than NHLF seeded at the same density (Figure 2). Thus, contractility can be directly linked to the disease state, opening a new approach for the characterization of fibroblast activation and as a potential evaluation method of novel small molecule efficacy. Notably, the capacity of fibroblasts to contract was maintained despite the decreased expression of *ACTA2* and *CTGF* compared to tissue culture plastic (Figure 3). Concurrently, these cells also reduce the expression of proliferation markers, indicating the loss of the classical *in vitro* myofibroblast state.

As we characterized the altered gene expression in fibroblasts seeded in these IPF hydrogels, we were struck by the inflammatory gene signature (Figure 3C). We have previously described the altered expression of an oxidative stress-induced cytokine axis when IPF fibroblasts are removed from the lung and seeded in 2D on a tissue culture dish(Rodriguez *et al*., 2018). In that work, we demonstrated the sensitivity to oxidative stress and the proinflammatory state of the IPF-F under a hydrogen peroxide challenge. We returned to that same data set and interrogated a series of genes that changed expression in IPF-F and NHLF when these cells were isolated from the lung and seeded on tissue culture plastic (Figure 4A). Interestingly, we observe that IPF-F seeded in the IPF hydrogels augment gene expression of several inflammatory cytokines.

Together, these data suggest that IPF-F in the 3D hydrogel system enter a proinflammatory phenotype and that this system potentially models an emerging role for fibroblasts as immune signaling hubs(Sundararaj *et al*., 2009; Österreicher *et al*., 2011; Kolahian *et al*., 2016; Moss *et al*., 2022; Xu *et al*., 2022).

To better explore the potential inflammatory signaling initiated by the reintroduction of fibroblasts to IPF ECM, we developed a series of co-culture experiments that aimed to model the crosstalk between fibroblasts and immune cells. Consistent with our observation of increased inflammatory expression in fibroblasts cultured in IPF hydrogels (Figures 3-4), we describe differential effects when comparing monocytes and macrophages exposed to fibroblast ECM conditioned media (Figures 6-7). We broadly defined our gene signatures as either proinflammatory or pro-resolution. Although this classification simplifies the complex nature of monocyte and macrophage biology, it allows us to identify intriguing differences between the monocyte and macrophage response to the conditioned media from fibroblast populations. Specifically, we observed increased cytokine expression in monocytes treated with conditioned media from fibroblasts grown in the IPF hydrogel. Independent of our proinflammatory or pro-resolution nomenclature, this observation suggests that cytokine expression within fibroblasts exposed to the fibrotic ECM is a potential signaling mechanism for the activation and/or recruitment of circulating monocytes. We also observed an increase in proliferation-associated gene expression, indicating that, once recruited to the site of injury, fibroblast crosstalk may also promote the expansion of the recruited monocyte population (Figure 8). The role of monocytes recruited to the IPF lung as biomarkers and drivers of disease progression has been a topic of great interest in recent years(Greiffo *et al*., 2016; Kolahian *et al*., 2016; Kreuter *et al*., 2021) and our model suggests that the fibrotic ECM itself may contribute a stimulus to initiate this crosstalk. Of particular interest, was the significant increase in expression of *CD86* and *IL10* in monocytes when exposed to IPF-F ECM conditioned media compared to the NHLF equivalent, suggesting that isolated IPF-F may retain a higher sensitivity to matrix cues (Figure 6B). IL-10, a ligand that has been implicated in monocyte-derived macrophage differentiation and IPF pathobiology(Millar, 2006; Foster *et al*., 2015; Greiffo *et al*., 2016; She *et al*., 2021) highlights that incoming monocytes exposed to signals from IPF-F fibroblasts may more rapidly differentiate into tissue specific macrophages. Furthermore, the differential increase in CD86, a surface marker known to be expressed in inflammatory-associated macrophages and a co-stimulatory molecule that stimulates T-cell responses(Zhang *et al*., 2018; Lis-López *et al*., 2021), supports this accelerated crosstalk. As a future scope of this work, additional studies applying omics technology are required to investigate the biological implications of these observations thoroughly.

The conditioned media challenge in macrophages had a similar effect with respect to in macrophages; however, there were some critical differences. The conditioned media derived from IPF-F and NHLF seeded in ECM hydrogels continued to increase the proinflammatory signature in macrophages. However, the genes listed as pro-resolution were either decreased or unaffected in macrophages, save for CCL24 (Figure 7A-B). This challenge broadly supports the hypothesis that fibroblasts seeded in IPF ECM are primed to secrete cytokines that drive an inflammatory monocyte and/or macrophage phenotype. The observation of increased expression of CCL24, a specific eotaxin recently identified as a crucial player in IPF biology and acts as a promising therapeutic target, underscores these findings.

Taken together, our findings suggest that there are properties inherent to the reported unique hydrogel system that result in a fibroblast with a less “classically active” but more inflammatory phenotype than those grown on tissue culture plastic. We see this inflammatory signature in the fibroblast and its crosstalk with immune cells, which implies that the fibrotic ECM, by its nature, contains inflammatory cues that contribute to this disease. This inflammatory induction property is attributable to the protein composition of the matrix and is a promising indication that fibrotic hydrogels recapitulate aspects of the *in vivo* lung environment that are not captured by tissue culture plastic. These data are also supported by previous findings from our group and others, describing disease-associated fibroblast alterations to their functional biology(Emblom-Callahan *et al*., 2010; Rodriguez *et al*., 2018). Fibrotic ECM hydrogels provide a potentially more reflective way of assessing fibroblasts and diseases associated with matrix remodeling. By developing materials for fibroblast culture that mimic physiological conditions, we hope to expand knowledge on novel pathways toward screening, discovery, and approval of more efficacious antifibrotic therapeutics.

## 5. AUTHOR CONTRIBUTIONS

LR, GG, JLM conceived and designed research; JFD, DM, JK, JAK, AKS, ML performed experiments and collected samples; JFD, DM, AKS, SD, LR, GG, JLM analyzed data and interpreted results; JFD generated figures and drafted the manuscript; JFD, AKS, CSK, SD, LR, GG, JLM revised the manuscript; all authors have reviewed and approved the final, submitted version of the manuscript and agreed to be listed as authors.

## 6. ACKNOWLEDGMENTS

The authors would like to thank all the patients who donated their tissue and allowed this work to be done. We also extend our gratitude to the clinical coordinators at Inova who reached out and obtained approval from all the patients. Finally, we thank Dr. Tom Huff, Tabitha King, Amy Carfagno, and Ammal H Altalhi, for their technical guidance.

## 7. FUNDING

This research was supported by the Virginia Commonwealth Health Research Board (grant #247-03-21).

## 8. DISCLOSURES

No conflicts of interest, financial or otherwise, are declared by the authors.

## Notes

### Competing Interest Statement

The authors have declared no competing interest.

### Summary of Updates

Additional Figure to the main text and updated supplemental tables to include reagents used in the additional figure.

