## Supplemental Data for "Pulmonary Matrix Derived Hydrogels from Patients with Idiopathic Pulmonary Fibrosis Induce a Proinflammatory State in Lung Fibroblasts *In Vitro*"

### SUPPLEMENTAL TABLES

| Supplemental Table 1: Antibodies | | | |
|---|---|---|---|
| **Antibody** | **Clone** | **Catalog Number** | **Manufacturer** |
| **PCNA** | PC10 | 2586S | Cell Signaling |
| **B-Actin** | 2D4H5 | 66009-1-Ig | Proteintech |

| Supplemental Table 2: Primer Sequences | | |
|---|---|---|
| **Target** | **Forward** | **Reverse** |
| 18S | GATGGGCGGGGAAAAATAG | GCGTGGATTCTGCATAATGGT |
| COL1A1 | GGACACAGAGGTTTCAGTGG | CCAGTAGCACCATCATTTCC |
| ACTA2 | AGTTACGAGTTGCCTGATGG | GAGGTCCTTCCTGATGTCAA |
| CTGF | CAGCATGGACGTTCGTCTG | CCAACCACGGTTTGGTCCTT |
| CCL20 | ATTGTGCGTCTCCTCAGTAAA | ACAAGTCCAGTGAGGCACAAA |
| CXCL2 | GAAAGCTTGTCTCAACCCCCG | TGGTCAGTTGGATTTGCCATTTT |
| CXCL3 | CCCAAACCGAAGTCATAGCCA | ACCCTGCAGGAAGTGTCAAT |
| CCL13 | GCCCAGTTTGGTTCTGAAGATGA | GCACTCAACGTCCCATCTACT |
| CXCL14 | AAGGGACCCAAGATCCGCTA | TCTTCGTAGACCCTGCGCTT |
| CXCR4 | GAGAAGCATGACGGACAAGTA | TGACAATACCAGGCAGGATAA |
| CD163 | GTCGCTCATCCCGTCAGTCATC | GCCGCTGTCTCTGTCTTCGC |
| CD86 | CACGGATGAGTGGGGTCATT | AGAGGAGCAGCACCAGAGA |
| IL6 | TTCGGTCCAGTTGCCTTCTCC | GTTGTTTTCTGCCAGTGCCTC |
| CCL8 | CTTGCCCTCCAAGATGAAGGT | CTGACCCATCTCTCCTTGGG |
| CCL7 | TCCAATTCTCATGTGGAAGCC | AGTCCTGGACCCACTTCTG |
| TNFA | GCTGCACTTTGGAGTGATCG | GCTTGAGGGTTTGCTACAACA |
| IL-1B | ATGATGGCTTATTACAGTGGCA | AAGCCCTTGCTGTAGTGGTG |
| CCL24 | CTGCAAGGACCCGAGCTATT | GATGACCACAGAGCCCGTAG |
| TOP2A | CACGACCGTCACCATGGAA | TGTTGTCCGCAGCATTAACTAGA |
| MKI67 | GGATCGTCCCAGTGGAAGAG | CAAACAAGCAGGTGCTGAGG |
| P16 | TGAGCTTTGGTTCTGCCATT | AGCTGTCGACTTCATGACAAG |
| IL10 | GGCACCCAGTCTGAGAACAG | GGCAACCCAGGTAACCCTTA |
| PCNA | GGCTCCATCCTCAAGAAGGTG | GGGACGAGTCCATGCTCTG |
| TGFB | CGGCCTTTCCTGCTTCTCA | ACTTCCAGCCGAGGTCCTT |

### SUPPLEMENTAL FIGURES

#### SUPPLEMENTAL FIGURE S1

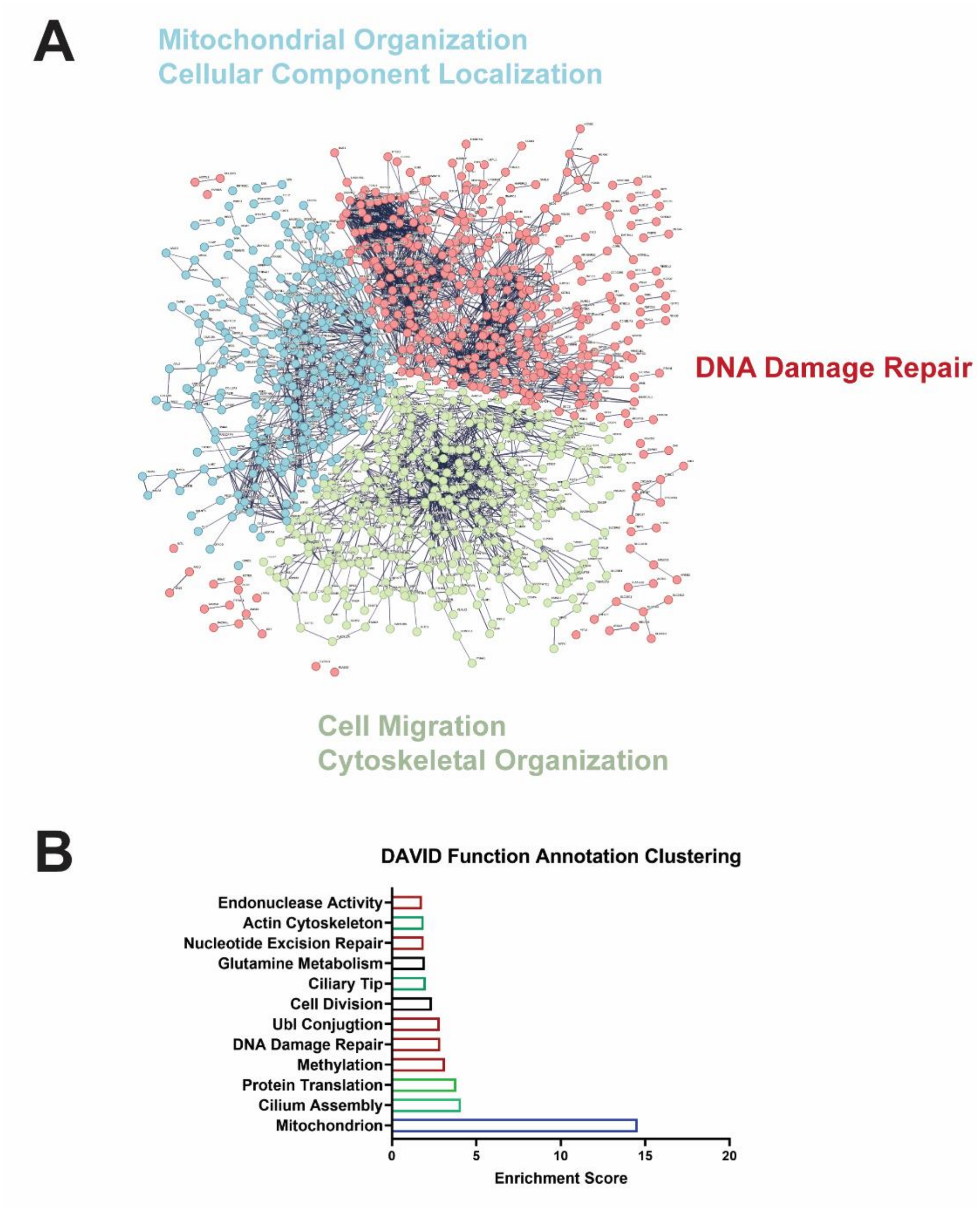


*Supplemental Figure S1*: (A) STRING DB analysis of genes upregulated during IPF fibroblast transition from i*n-vivo* environment to tissue culture plastic reveals three broad pathways that are upregulated. (B) DAVID Functional Annotation of this gene subset confirms STRING DB analysis and further defines complementary pathways increased in IPF-F through expansion on tissue culture plastic.

**SUPPLEMENTAL FIGURE S2**

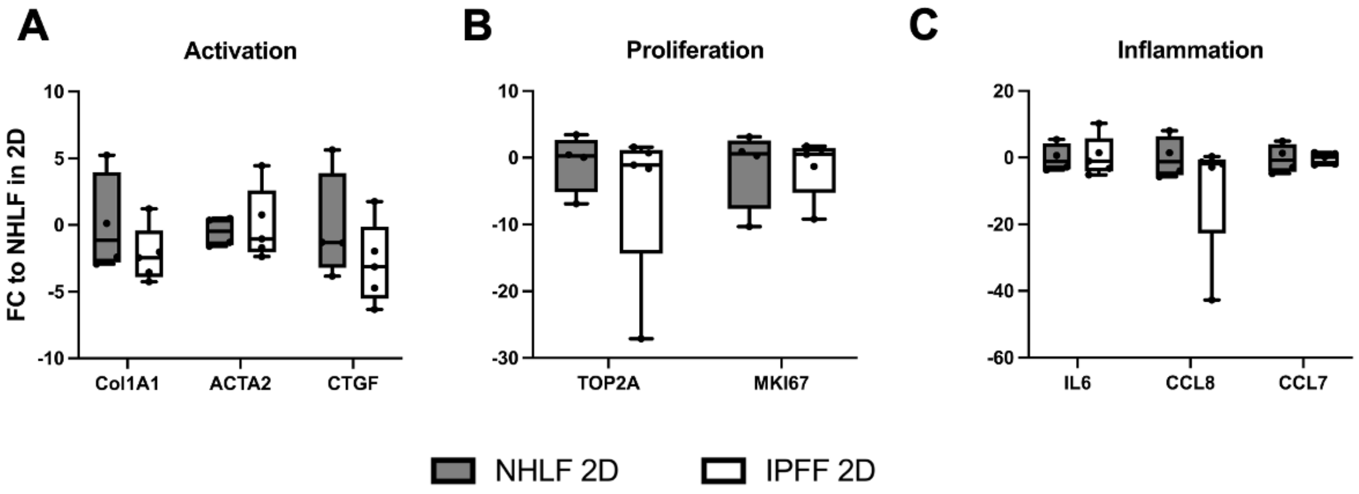


*Supplemental Figure S2: Fold Change of Baseline Gene Expression of IPF-F and NHLF Seeded in 2D Tissue Culture Plastic Normalized to NHLF in 2D:* (A) Gene expression of fibroblast activation markers: *Col1a1, ACTA2,* and *CTGF*. (B) Gene expression of markers associated with cell proliferation: *(*C) *TOP2A* and *MKI67*Gene expression of ligands associated with inflammation: *IL6, CCL8, and CCL7*. *: $p<0.05$.

**SUPPLEMENTAL FIGURE S3**

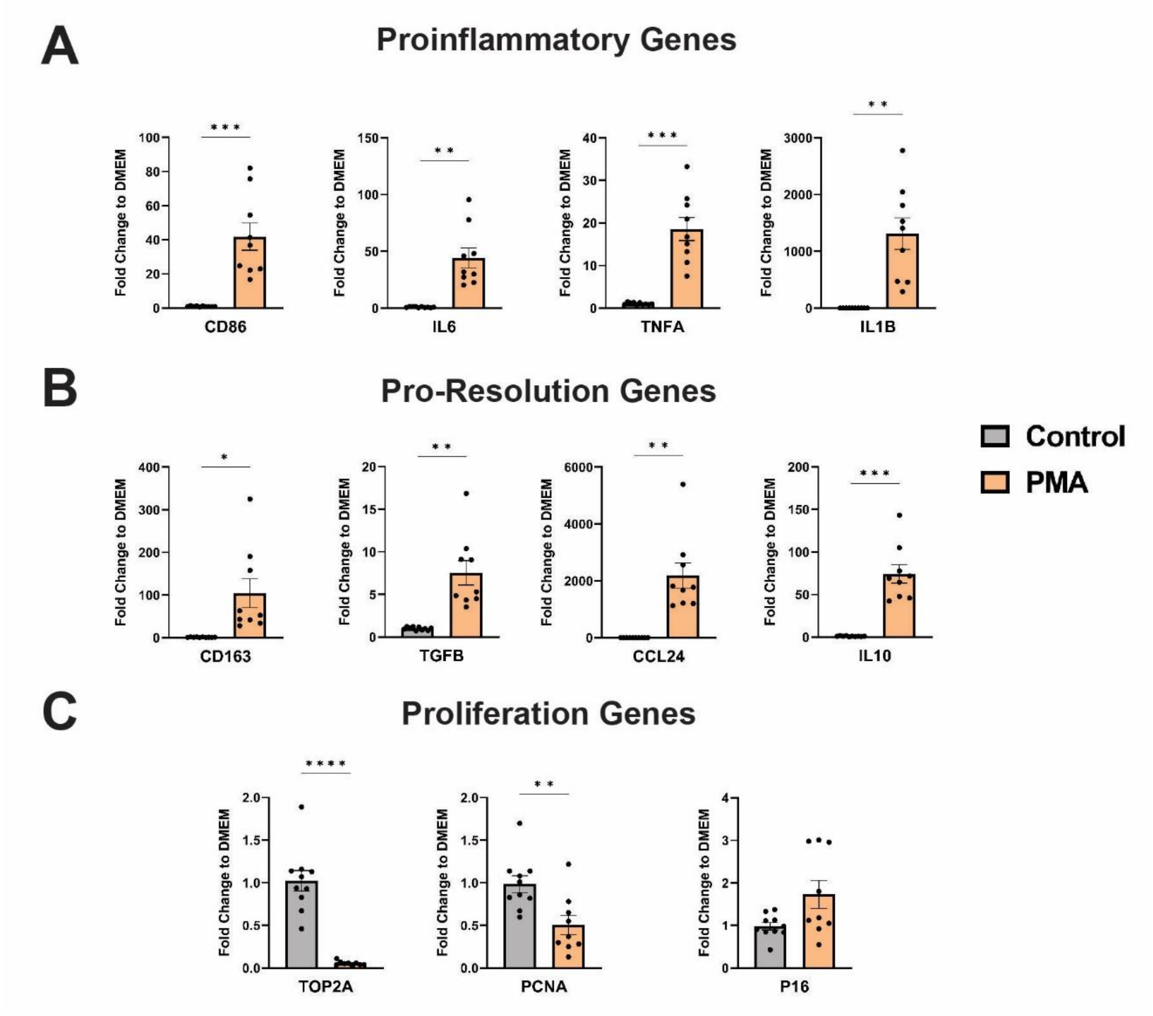


*Supplemental Figure S3*: Gene expression in monocytes after 48-hour PMA challenge. (A-B) The 48-hour PMA challenge induces the expression of genes that we have categorized as proinflammatory and pro-resolution. (C) Expression of proliferation-associated genes is inhibited after a 48-hour PMA challenge with a concurrent increase in cell cycle checkpoint inhibitor p16.
